# mRNA-1273 vaccines adapted to JN.1 or KP.2 elicit cross-neutralizing responses against the JN.1 sublineages of SARS-CoV-2 in mice

**DOI:** 10.1101/2024.12.24.629478

**Authors:** Diana Wing Lee, Arshan Nasir, Sayda Elbashir, Hardik Jani, Tessa Speidel, Amy Gorrie, Daniela Montes Berrueta, Philippa Martin, Swan Tan, Yixuan Jacob Hou, Kath Hardcastle, Darin Edwards, Kai Wu, Andrea Carfi, Yadunanda Budigi

## Abstract

The continued diversification of SARS-CoV-2 omicron lineage has given rise to the JN.1 variant and descendant strains (KP.2, KP.3, and XEC) that have prolonged the JN.1 infection wave. JN.1 and KP.2 show decreased susceptibility to neutralization sera in recipients of XBB.1.5 vaccine boosters, supporting the recent authorization of JN.1- and KP.2-matched mRNA vaccines in the United States, Europe, and other regions. We evaluated the immunogenicity of two updated monovalent variant-containing formulations of mRNA-1273 vaccines encoding the spike protein of the omicron subvariants JN.1 (mRNA-1273.167) and KP.2 (mRNA-1273.712) as compared with the monovalent XBB.1.5 vaccine (mRNA-1273.815). The vaccines were administered either as a two-dose primary series in naive mice or as a booster (third) dose in mice previously immunized with two-dose primary series of mRNA-1273 (ancestral strain). The neutralizing antibody response elicited by these vaccines against JN.1 subvariants (KP.3 and LA.2) and the recombinant strain (XEC), which achieved dominance in the United States during late 2024, was evaluated. Primary series immunization with either JN.1- or KP.2-matched vaccine elicited robust neutralizing antibody titers against the matched strains and effectively cross-neutralized KP.3, LA.2, and XEC, but not the antigenically distant XBB.1.5. Similarly, JN.1- and KP.2-matched vaccines administered as a booster (third) dose increased titers against the corresponding strains and JN.1-related subvariants, but not against XBB.1.5. These data suggest these strains are antigenically similar with relatively few spike differences between JN.1 and KP.2/JN.1-related subvariants. Our results demonstrate the potency of JN.1- and KP.2-containing mRNA-1273 vaccines in neutralizing the matched variants and their utility in cross-neutralizing JN.1-related subvariants KP.3, LA.2, and XEC. Taken together, these data suggest that the licensed JN.1 and KP.2 mRNA vaccines are likely to continue to protect against the emerging strains as the JN.1 lineage further evolves.

## 1. Introduction

The SARS-CoV-2 virus is constantly evolving by accumulating mutations in the spike gene and/or through the recombination of co-circulating strains [1]. The emerging SARS-CoV-2 variants, including the most recent and highly diversified omicron lineages, contain mutations that contribute to escape from both natural and vaccine-induced humoral immunity [1,2]. This necessitates updates to the antigenic composition of COVID-19 vaccines to maintain protection against the circulating variants [3,4]. In response to the evolving pandemic, variant-containing mRNA-1273 booster vaccines that encode the spike protein of the SARS-CoV-2 variant lineages were authorized/approved in 2022, 2023, and 2024, including a bivalent mRNA-1273 vaccine (mRNA-1273.222) encoding the ancestral SARS-CoV-2 and omicron BA.4/BA.5 (2022-2023 formula); a monovalent mRNA-1273 vaccine (mRNA-1273.815) encoding the omicron sublineage XBB.1.5 (2023-2024 formula); and monovalent mRNA-1273 vaccines encoding the omicron sublineages KP.2 or JN.1, which were approved in the United States, European Union, and Canada for the 2024-2025 season [4–7].

JN.1 is a descendant of omicron BA.2.86, which harbors >30 mutations in the spike protein compared with the parental BA.2 lineage [8,9]. JN.1 was designated a variant of interest (VOI) by the World Health Organization (WHO) as of December 2023 [10], due to its high potential for immune evasion and increasing global prevalence [8,11]. Compared with BA.2.86, JN.1 contains an additional mutation in the receptor-binding domain (RBD) at L455S, which contributes to its increased transmissibility and enhanced immune escape [8,12]. The effectiveness of previously authorized XBB.1.5-adapted mRNA vaccines against infection, hospitalization, and death was significantly reduced in the setting of JN.1 variant predominance [2]. In virus neutralization assays, significantly lower titers against JN.1 compared with BA.2.86 were observed in sera of individuals vaccinated with monovalent XBB.1.5 vaccine [8], while sera from individuals who experienced breakthrough JN.1 infection exhibited high neutralization titers against JN.1, BA.2.86, and BA.2, and lower cross-neutralization against XBB descendant lineages (XBB.1, EG.5.1, and HK.3), supporting the antigenic distance between XBB and JN.1 [12]. These data suggested that cross-neutralization of JN.1 by XBB.1.5 vaccines in human sera might be limited [8,12] and supported the need for development of updated vaccines matched to emerging strains in the JN.1 lineage [5]. Accordingly, WHO Technical Advisory on COVID-19 Vaccine Composition and multiple public health agencies recommended the use of updated COVID-19 vaccines (2024-2025 season) targeting the JN.1 family of omicron subvariants (JN.1 and KP.2) [3,4,13]. Recommendations were based on the virus epidemiology data at that time, performance of approved vaccines against circulating SARS-CoV-2 variants, as well as evidence of preclinical activity of updated vaccines by vaccine manufacturers [13,14].

Compared with JN.1, the KP.2 variant contains three additional spike mutations, including two mutations in the RBD (R346T and F456L) [15,16]; was reported to have a higher relative effective reproduction number (R_e_), suggesting higher viral fitness [15]; and demonstrated enhanced evasion of serum antibodies, likely conferred by the acquisition of F456L [17]. KP.2 and its antigenically related variants KP.3, KP.3.1.1, LB.1, and XEC were classified as variants under monitoring by WHO as of September 2024 [10]. XEC in particular is emerging in several parts of the world, including in the United States [18], where it is now the most commonly sequenced strain [19]. XEC is a recombinant strain involving genetic elements from two JN.1 subvariants (KP.3.3 and KS.1.1) with a recombination breakpoint inside the spike gene [18,20]. The small number of spike differences in the currently circulating strains compared with approved vaccine compositions suggest they are likely to be antigenically similar, and therefore cross-neutralized by neutralizing sera. However, this remains to be fully corroborated by clinical studies from recipients of the updated vaccines.

We evaluated the neutralizing antibody (nAb) responses of the monovalent SARS-CoV-2 vaccine encoding variants JN.1 (mRNA-1273.167) and KP.2 (mRNA-1273.712) administered as a two-dose primary series in naive mice, or as a booster (third) dose in mice previously immunized with the original mRNA-1273. A further objective was assessing cross-neutralization potential of these vaccines against additional variants in the JN.1 lineage, including KP.3 and LA.2, as well as the recombinant strain (XEC) that achieved dominance in the United States as of late 2024 [18,19].

## 2. Methods

### 2.1. Sequence analysis

Genetic sequences for SARS-CoV-2 were downloaded from the GISAID database [21] and aligned to the ancestral strain (GenBank Accession Id: MN908947) via the Nextclade pipeline (v. 3.8.2; https://clades.nextstrain.org) [22] and in-house Python scripts (v. 3.12.2) for quality control, clade and variant assignment, and mutation calling relative to the ancestral strain.

### 2.2. mRNA vaccines

The monovalent mRNA-1273.167 vaccine evaluated in this study contained a single mRNA encoding the SARS-CoV-2 spike protein of the omicron SARS-CoV-2 subvariant JN.1 with two proline substitutions within the heptad repeat 1 domain (S-2P) antigen. The monovalent mRNA-1273.712 vaccine contained a single mRNA encoding the SARS-CoV-2 S-2P antigen of the JN.1 subvariant KP.2. A monovalent mRNA-1273 vaccine, which was administered as a primary series (doses 1 and 2) in the booster study, contained a single mRNA encoding the SARS-CoV-2 S-2P antigen of the ancestral SARS-CoV-2 strain. In all studies, a monovalent mRNA-1273.815 vaccine, which contained a single mRNA encoding the SARS-CoV-2 S-2P antigen of the omicron subvariant XBB.1.5, served as the active control. mRNAs were encapsulated in a lipid nanoparticle (LNP) through a modified ethanol-drop nanoprecipitation process [23].

### 2.3. Mouse model

Female BALB/c mice (6 to 8 weeks old; Charles River Laboratories) were used. Animal experiments were carried out with the approval of the ModernaTX, Inc. Institutional Animal Care and Use Committee (IACUC Protocol 22-02-00) and in compliance with all applicable local and federal regulations including the National Research Council’s Guide for the Care and Use of Laboratory Animals. mRNA formulations were diluted in 50 µL of 1X phosphate-buffered saline (PBS); control mice received PBS. The groups in the study were not blinded.

### 2.4. Study design and treatment groups

In the primary series studies, mice were immunized into the same hind leg 3 weeks apart (days 1 and 22) with two intramuscular (IM) injections of 1 µg of test articles (mRNA-1273.815, mRNA-1273.167, or mRNA-1273.712) or PBS (n=7-8 per group in the experiment evaluating mRNA-1273.167 and n=8 in the experiment evaluating mRNA-1273.712). Blood was collected from all animals on day 21 (before the administration of dose 2) and day 36 (2 weeks after administration of the primary series [dose 2]). In the booster studies, mice (n=8 per group) were immunized into the same hind leg 3 weeks apart (days 1 and 22) with two IM injections of mRNA-1273 (0.5 μg) and were then boosted 19 days or 28 days later with a single IM injection into the same hind leg (1.0 μg) of the test articles (mRNA-1273.712, mRNA-1273.167, or mRNA-1273.815). Blood samples were collected before administration of dose 3 (day 41 in the first booster study and day 51 in the second booster study) and 2 weeks after dose 3 (day 56 in the first booster study and day 69 in the second booster study).

### 2.5. Assays

Serum samples were analyzed for nAb responses via vesicular stomatitis virus (VSV)-based pseudovirus neutralization assay (PsVNA). Codon-optimized full-length spike genes (XBB.1.5, JN.1, and KP.2) were cloned into a pCAGGS vector. The recombinant VSVΔG-based SARS-CoV-2 pseudovirus was generated by transfecting BHK-21/WI-2 cells with the spike expression plasmid and subsequently infected by VSVΔG-firefly-luciferase as described previously [26]. For neutralization assay, A549-hACE2-TMPRSS2 cells were used as target cells. Mouse serum samples were heat-inactivated for 45 minutes at 56°C, and serial dilutions were made in Dulbecco’s Modified Eagle Medium supplemented with 10% fetal bovine serum. The diluted serum samples or culture medium (serving as virus-only control) were mixed with VSVΔG-based SARS-CoV-2 pseudovirus and incubated at 37°C for 45 minutes. The inoculum virus or virus-serum mix was subsequently used to infect A549-hACE2-TMPRSS2 cells for 18 hours at 37°C. At 18 hours after infection, equal volume of One-Glo reagent (Promega) was added to the culture medium for readout; the percentage of neutralization was calculated based on relative luminescence units (RLU) of the virus-only control and was analyzed using four-parameter logistic curve (Prism v.9).

Serum samples were analyzed for binding antibody responses by the enzyme-linked immunosorbent assay (ELISA; **Supplementary Methods**).

### 2.6. Statistical analysis

The neutralization titers were summarized descriptively. Differences in serum nAb responses against VSV-based PsVNA for XBB.1.5, JN.1, KP.2, KP.3, LA.2 and XEC strains within each treatment group and across treatment groups were described as fold changes or geometric mean values. All statistical analyses were performed using a two-tailed Wilcoxon paired samples test. Data are presented as geometric mean titers (GMTs) ± 95% confidence intervals.

## 3. Results

### 3.1. Ongoing antigenic evolution in SARS-COV-2 strains

Emergence of the highly-mutated BA.2.86 variant was followed by the appearance of JN.1 strain, containing an additional RBD mutation over BA.2.86 [8,9,16]. JN.1 quickly displaced the XBB family of subvariants dominant at that time and became globally dominant by early 2024 [11]. Since then, a number of JN.1 subvariants have emerged either via incremental evolution in the JN.1 spike protein (eg, KP.2, KP.3, and LA.2) or a recombination event involving JN.1 subvariants (eg, XEC) [15,18,27,28], which have prolonged the JN.1 infection wave. The JN.1 subvariants are characterized by a small number of mutations in the N-terminal domain (NTD) and RBD (**Figure 1A**), the latter being the main target for neutralizing antibodies [29]. KP.2 and KP.3 caused infection waves from summer and early fall 2024 [30,31], and are now challenged by the rapid spread of XEC, especially in Europe and the United States [18]. However, unlike the significant number of amino acid differences observed between JN.1 and XBB.1.5 (2023-2024 vaccine strain), there are relatively fewer spike differences between JN.1 and KP.2 or emerging JN.1 subvariants (**Figure 1B**), suggesting that they are likely to be antigenically related in the context of neutralizing responses.

**Figure 1.**
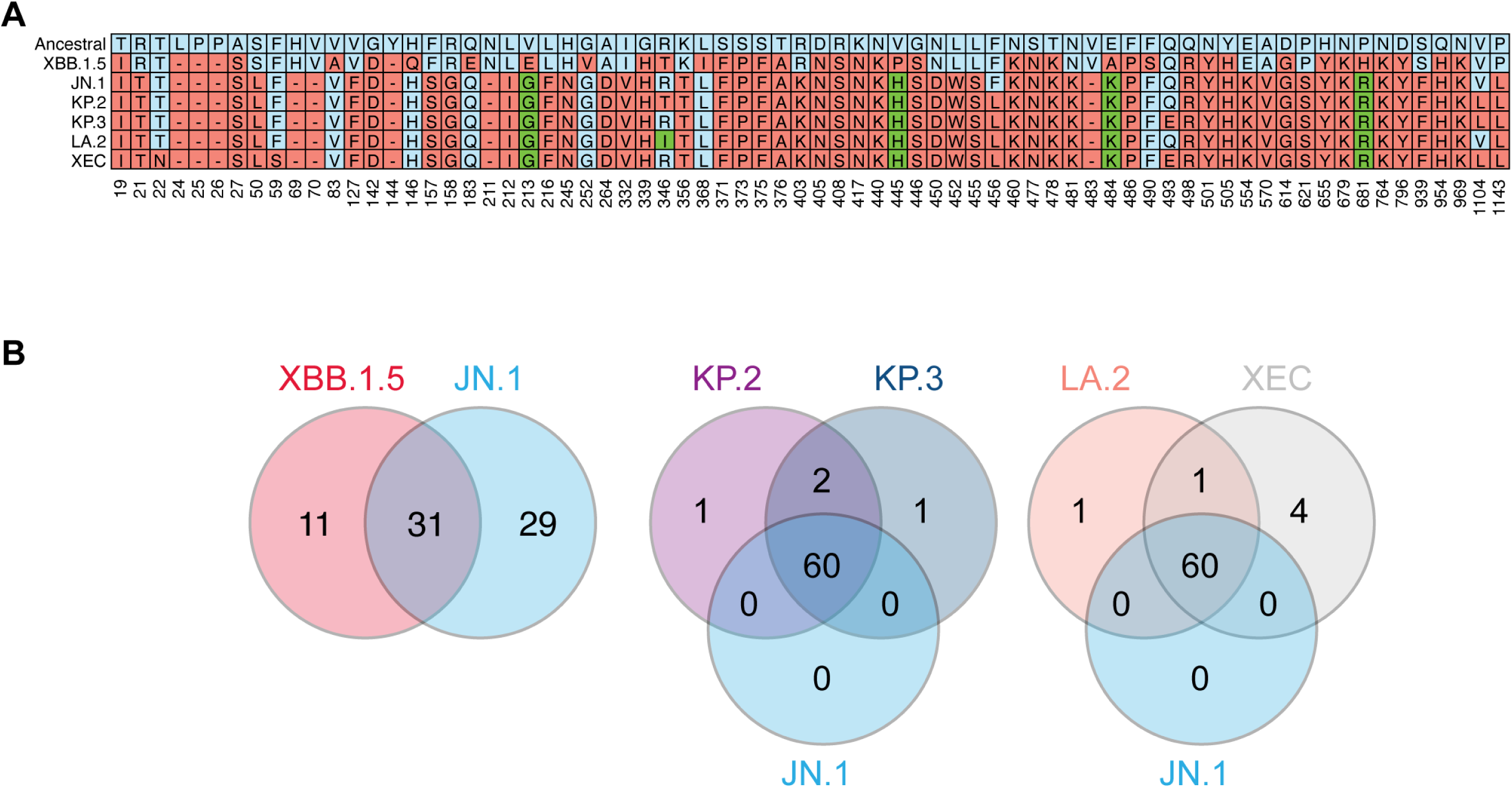
Spike amino acid variations in the variants analyzed. **(A)** The heatmap highlights spike mutations that have emerged in variants XBB.1.5, JN.1, KP.2, KP.3, LA.2, and XEC, analyzed in this study. The x-axis indicates amino acid position relative to the ancestral SARS-CoV-2 strain. Mutations are indicated in red; sites that have mutated more than once (213, 346, 445, 484, and 681) are indicated in green. JN.1 and related subvariants also encode a 4 amino acid insertion (MPLF at position 16) in the spike NTD that is not shown. **(B)** Relatively few changes in the RBD domain are observed for the recent variants KP.3, LA.2, and XEC relative to JN.1 or KP.2. Venn diagrams show the number of unique and shared spike mutations among analyzed variants relative to the ancestral strain. Different substitutions at same sites (eg, 446) and contiguous deletions (eg, 24-27) were counted separately. Insertion at position 16 (16MPLF) was not counted.

### 3.2. Neutralizing antibody responses after JN.1 and KP.2 primary series vaccination in mice

To evaluate the immunogenicity of a two-dose primary series, naive BALB/c mice were immunized on day 1 and day 22 in two separate studies with either a monovalent JN.1 vaccine (mRNA-1273.167; 1 µg) or a monovalent KP.2 vaccine (mRNA-1273.712; 1 µg). In each study, separate groups of mice were immunized with mRNA-1273.815 (XBB.1.5 vaccine; 1 µg) or PBS. Serum samples were collected on day 36 (14 days post-primary series vaccination) and analyzed for nAb responses against JN.1, KP.2, KP.3, LA.2, XEC, and XBB.1.5.

At 14 days post-primary series vaccination, mice that received mRNA-1273.167 exhibited high serum nAb titers against JN.1 (GMT, 9758). These animals also exhibited high nAb titers against JN.1-related subvariants KP.2, KP.3, LA.2, and XEC (GMT range, 3609-7666), but low nAb titers against XBB.1.5 (GMT, 47) (**Figure 2A**). The nAb titers elicited by mRNA-1273.167 against JN.1 subvariants relative to titers against JN.1 ranged from 1.3- to 2.7-fold lower compared with nAb titers against XBB.1.5, which were >200 fold lower. These data indicate robust cross-neutralization of all JN.1 sublineages by mRNA-1273.167.

**Figure 2.**
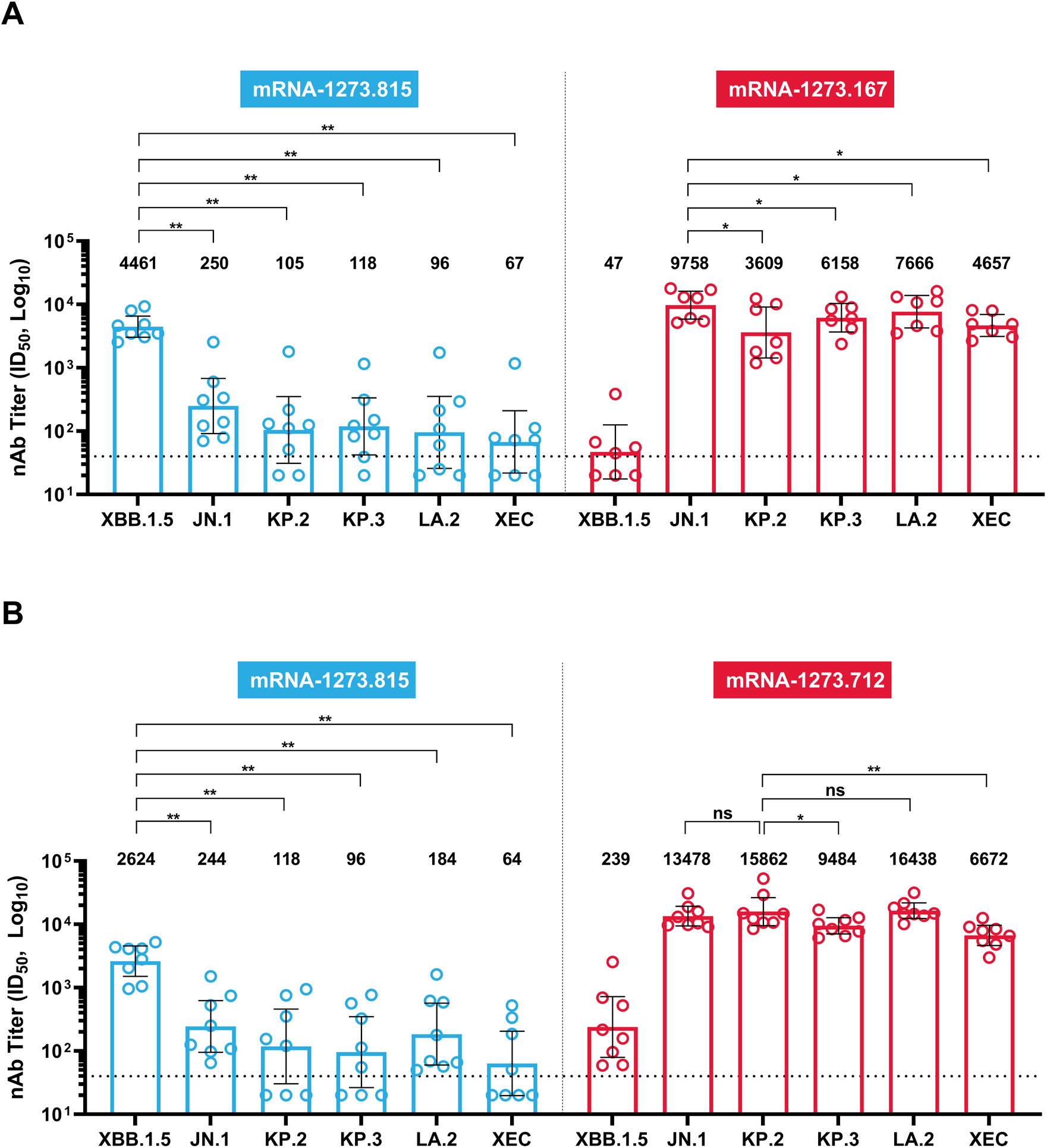
Neutralizing antibody responses against JN.1-related subvariants (KP.2, KP.3, LA.2, XEC) and XBB.1.5 14 days after primary series vaccination (1 µg) in BALB/c mice with **(A)** mRNA-1273.167 (JN.1 variant vaccine)^a^ and **(B)** mRNA-1273.712 (KP.2 variant vaccine), compared with mRNA-1273.815 (XBB.1.5 variant vaccine). The numbers and bars represent GMTs, and whiskers represent the 95% CI for day 36 (14 days after primary series). The dotted line indicates the LOD of 1 in 40 for the neutralization assay. Statistical differences were defined using a two-tailed Wilcoxon paired samples test; *P*<0.05 = *; *P*<0.01 = **; *P*<0.001 = ***; ns, non-significant. ^a^One animal died in the mRNA-1273.167 group on Day 21 due to the bleeding procedure; no sample was obtained from this animal for analysis. *Abbreviations*: CI, confidence interval; GMT, geometric mean titer; ID_50_, inhibitory dilution 50%; LOD, limit of detection; mRNA, messenger RNA; nAb, neutralizing antibody.

In a similar study, mice vaccinated with a primary series of mRNA-1273.712 (KP.2 vaccine) showed high serum nAb titers against KP.2 (GMT, 15,862) and related subvariants (GMT range, 6672-16,438), but poor cross-neutralization against XBB.1.5 (GMT, 239) (**Figure 2B**). mRNA-1273.712 elicited comparable nAb titers against JN.1, KP.2, and LA.2 (GMT, 13,478, 15,862, and 16,438, respectively) and slightly lower (~1.7- and ~2.4-fold, respectively) titers against KP.3 and XEC; titers elicited against XBB.1.5 were >65-fold lower.

In both studies, sera from mice vaccinated with primary series of mRNA-1273.815 exhibited high serum neutralization titers against XBB.1.5 and low titers against JN.1 and KP.2-related subvariants (**Figure 2**), suggesting very low levels of cross-neutralization. No nAb titers were detectable in the PBS control group (data not shown). Neutralizing antibody titers against JN.1 subvariants elicited by updated JN.1 and KP.2 vaccines were ~40- to 130-fold higher than those elicited by the XBB.1.5 vaccine, underscoring the antigenic distance between these strains. These differences were not related to differences in immunogenicity of these antigens, as they elicited comparable spike binding immunoglobulin G (IgG) titers (**Supplementary Figure 1**). The data indicate that mRNA-1273.167 and mRNA-1273.712 were immunogenic and elicited high neutralization titers that cross-neutralized a wide breadth of JN.1 sublineage strains, including the recently circulating XEC.

### 3.3. Neutralizing antibody responses after JN.1 and KP.2 booster vaccination in imprinted mice

With immunity from SARS-CoV-2 vaccines and infection being widespread in the global population [5], it is important to assess performance of variant vaccines in a background of immune imprinting. While it is not feasible to fully recapitulate the human vaccine/virus exposure history, we have previously tested the boosting potential of variant vaccines in animals administered a primary series vaccination to inform on vaccine performance [32]. Using this approach in two separate studies, we tested the immunogenicity of a booster (third) dose of JN.1 vaccine (mRNA-1273.167; 1 µg) or KP.2 vaccine (mRNA-1273.712; 1 µg) in BALB/c mice administered approximately 2-4 weeks after the second dose of a two-dose primary series immunization with mRNA-1273 (ancestral strain; 0.5 µg).

In the first study, we compared groups administered mRNA-1273.167, mRNA-1273.712, or mRNA-1273.815 ~3 weeks (19 days) after the second dose of mRNA-1273. Blood samples were collected for evaluation of nAb responses before the administration of the booster dose (day 41) and 14 days after the booster dose (day 56). To assess the impact of duration between second and third dose, we also conducted a second study where the duration between second primary series dose (mRNA-1273) and booster (third) dose of mRNA-1273.167 or mRNA-1273.815 was extended to ~4 weeks. Blood samples were collected for evaluation of nAb responses before the administration of the booster dose (day 51) and approximately 14 days after the booster dose (day 69). We measured nAb responses against JN.1, KP.2, KP.3, LA.2, and XBB.1.5 prior to and 2 weeks after the booster dose.

At 19 days after vaccination with mRNA-1273 (day 41), pre-boost nAb titers were generally below or close to the detection limit against XBB.1.5, JN.1, and KP.2/related subvariants (**Figure 3A**). At 14 days after booster (day 56), sera from mice boosted with mRNA-1273.167 showed high nAb titers against JN.1 (GMT, 120) and subvariants KP.2, KP.3, and LA.2 (GMT range, 62-76), while neutralization titers against XBB.1.5 remained close to the detection limit (**Figure 3A**). Similarly, at 14 days following a booster dose of mRNA-1273.712 (day 56), a significant boost of neutralization titers against variants JN.1, KP.2, KP.3, and LA.2 was observed (GMT range, 109-288), while nAb titers against XBB.1.5 remained below the detection limit (**Figure 3A**). By contrast, sera from mice boosted with mRNA-1273.815 (XBB.1.5 vaccine) showed increased nAb titers by ~3 fold relative to pre-boost levels against XBB.1.5, but little to no increase in titers against JN.1, KP.2, KP.3, or LA.2 variants (**Figure 3A**). All groups of mice showed comparable spike S2-P binding IgG titers (**Supplementary Figure 2**).

**Figure 3.**
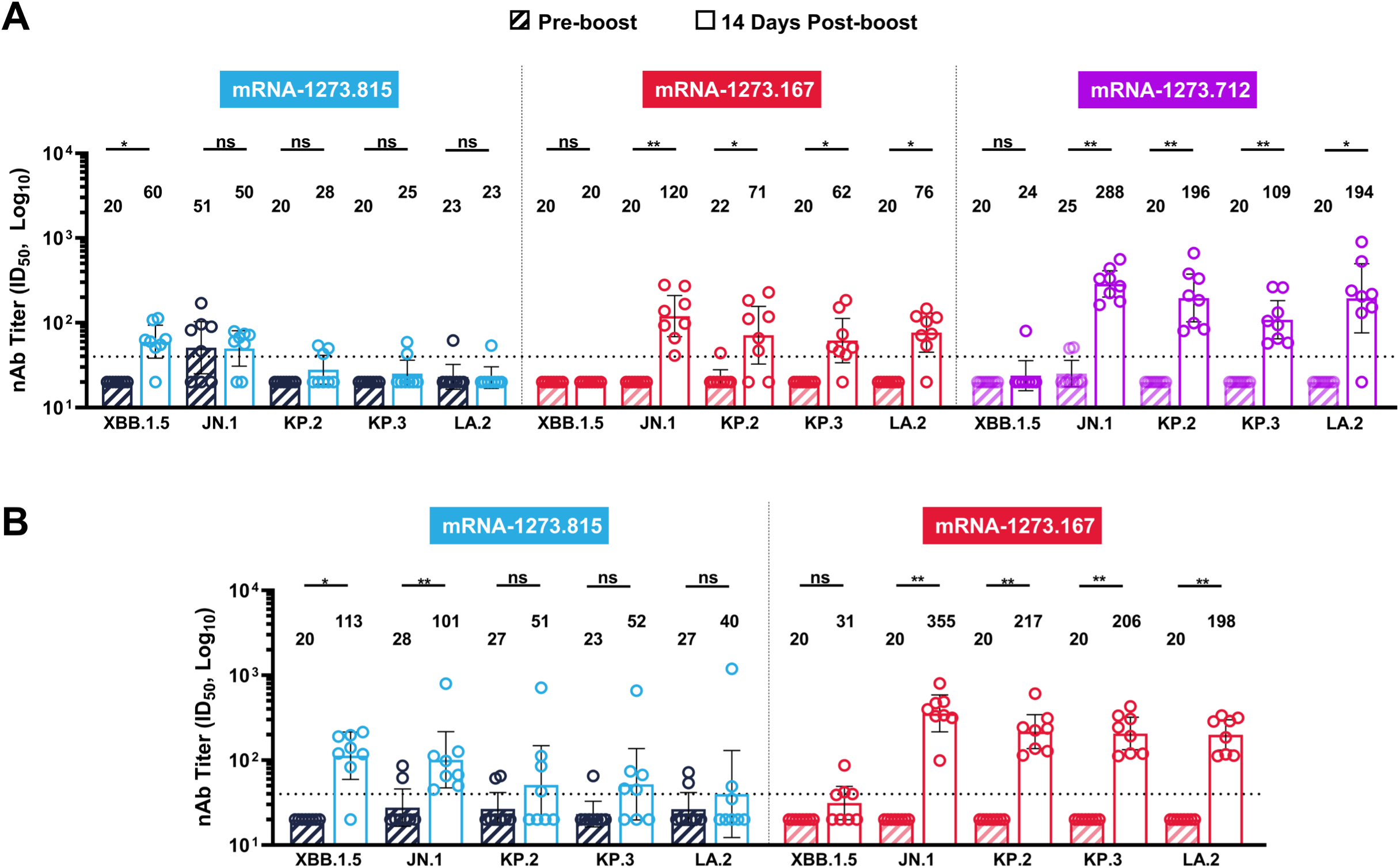
Boosting effect of mRNA-1273 variant vaccines in imprinted BALB/c mice. **(A)** Neutralization titers at 14 days post booster (third) dose of mRNA-1273.167 (JN.1 variant vaccine), mRNA-1273.712 (KP.2 variant vaccine), or mRNA-1273.815 (XBB.1.5 variant vaccine) administered 19 days after the second dose of a two-dose primary series with mRNA-1273 (ancestral strain). **(B)** Neutralization titers at 14 days post booster (third) dose of mRNA-1273.167 or mRNA-1273.815 administered 28 days after the second dose of a two-dose primary series with mRNA-1273 (ancestral strain). The numbers and bars represent GMTs, and whiskers represent the 95% CI for pre-boost day 51 (4 weeks after primary series) and post-boost day 69 (2 weeks after booster [third] dose). The dotted line indicates the LOD of 1 in 40 for the neutralization assay. Statistical differences were defined using a two-tailed Wilcoxon paired samples test; *P*<0.05 = *; *P*<0.01 = **; *p*<0.001 = ***; ns, non-significant. *Abbreviations*: CI, confidence interval; GMT, geometric mean titer; ID_50_, inhibitory dilution 50%; LOD, limit of detection; mRNA, messenger RNA; nAb, neutralizing antibody.

In a second study we increased the interval between the second dose (mRNA-1273) and a booster (third dose) of mRNA-1273.167, which was administered 4 weeks after the second dose. In these mice, high nAb titers (>10-fold over pre-boost titers) were observed against JN.1, KP.2, KP.3, and LA.2 (GMT range, 198-355), while neutralization titers against XBB.1.5 remained close to the detection limit (**Figure 3B**). The post-boost titers observed in this study were higher than those in the previous study with a shorter dose interval (19 days) between the second and third doses. Notwithstanding the increased dose interval, while mRNA-1273.815 boosted nAb titers against XBB.1.5 by ~5-fold, there was little to no increase in nAb titers against JN.1, KP.2, KP.3, and LA.2 variants (**Figure 3B**). No nAb titers were detectable in the PBS control group (data not shown). Overall, the data indicate that updated mRNA-1273 vaccines matched to JN.1 lineage can boost nAb titers that effectively cross-neutralize JN.1 sublineage strains, even in the background of immune imprinting from previous exposure to an antigen encoded by the mRNA-1273 vaccine.

## 4. Discussion

Detection and genetic characterization of antigenically and epidemiologically relevant variants and routine evaluation of updated vaccine candidates is critical to maintain neutralization responses and provide continued protection against infection/severe disease caused by SARS-CoV-2 [5,33]. These require continuous monitoring of emerging variants, variant classification based on the presence of immune-evading mutations, and evaluation of updated vaccine compositions matched to these variants in preparation for authorization and deployment [5,33]. Mouse models have been informative for preclinical assessment of mRNA-LNP vaccines for evaluating inherent immunogenicity of candidate antigens and to assess immune escape [32,34,35]. Such models are key enablers of rapid development of updated vaccines in time for deployment as periodic boosters. Importantly, the overall conclusions drawn from mouse studies showing increased breadth of coverage against VOCs or VOIs after booster doses of variant-adapted mRNA-1273 vaccine are aligned with clinical observations [36,37].

We evaluated the immunogenicity of two formulations of monovalent variant-containing mRNA-1273 vaccines, one encoding the spike protein of the SARS-CoV-2 JN.1 variant (mRNA-1273.167) and another encoding the spike protein of the KP.2 variant (mRNA-1273.712), licensed for use in the United States, Europe, Japan and other regions of the world [38]. We tested these vaccines in two mouse models administered either as (1) a two-dose primary series in naive mice to assess the inherent immunogenicity in an immune naive background, or (2) a booster (third) dose in mice previously immunized with two doses of mRNA-1273 to evaluate immune responses in a background of immune imprinting with an antigenically distant strain. We have extensively used such studies to assess immunogenicity of our vaccine candidates [32,34,35], and, while such models cannot recapitulate the complex immune history of humans or the specific extent of response (eg, antibody titers), they serve as useful surrogates to assess gross responses. As expected, primary-series immunization with either JN.1 or KP.2 mRNA-1273 vaccine (mRNA-1273.167, mRNA-1273.712) effectively elicited robust nAb titers against the matched (JN.1, KP.2) strains, which did not cross-neutralize the XBB.1.5 strain, supporting the antigenic distance between these strains. The sera from these mice also showed robust cross-neutralization against other JN.1 sublineage strains KP.3, LA.2, and XEC. Following primary series immunization with JN.1 vaccine, the differences in neutralization against these various strains, as compared with JN.1, were generally within ~2-fold lower range, with the exception of KP.2, which was ~2.7 fold lower. This likely reflects the impact of the three additional mutations in KP.2, including two mutations in the RBD (F456L, R346T), which are not found in parental JN.1 [15,39]. Although the KP.3 variant lacks the spike protein substitution R346T present in KP.2, it contains the Q493E mutation on the RBD, which was shown to primarily affect ACE2-binding affinity and viral infectivity and has minimal impact on serum antibody neutralization [17,27,39]; this may explain, in part, numerically higher nAb titers against KP.3 relative to KP.2 following primary series vaccination with the JN.1 vaccine. By contrast, the XEC variant contains two additional mutations in the spike protein (T22N, F59S) compared with KP.3, which were acquired through recombination of KS.1.1 and KP.3.3 and were shown to contribute to increased virus infectivity and higher immune evasion of XEC [18]. T22N, in particular, introduces a new N-linked glycosylation motif in the XEC spike and likely has conformational effects in the spike protein [40] that might explain reduced neutralization of XEC compared with JN.1 as well as numerically lower neutralization relative to KP.3 after immunization with the JN.1 vaccine. It is noteworthy that the naive mouse background is especially sensitive to the incremental impact of such point mutations on immune evasion, which is in contrast with the more complex human immune history, where significant cross-reactive responses are likely to prevail and further limit immune escape [41,42].

We also evaluated immune responses in mice previously vaccinated with primary series of mRNA-1273 to assess impact of immune imprinting on responses with the expectation that imprinting effects would lead to a repeated boost of ancestral responses and a smaller boost of variant-specific response. Although we did not evaluate titers against ancestral strain, the variant vaccines matched to JN.1 or KP.2 boosted nAb titers against the corresponding strains and related JN.1 sublineages, but not against the more distant strain XBB.1.5. While heterologous variant boosting in previously exposed (imprinted) individuals has been suggested to drive cross-neutralizing responses [43], it is likely that the lack of boost in nAb titers against XBB.1.5 observed in our study is a reflection of the significant antigenic distance between not only the XBB.1.5 and JN.1 lineages, but also between the ancestral strain and XBB.1.5.

These findings are in keeping with recent data showing robust neutralization of JN.1 and BA.2.86 by sera from individuals after JN.1 breakthrough infection, with significantly reduced nAb titers against XBB.1 pseudovirus [12]. The antigenic mapping based on all the breakthrough serum neutralization data in this cohort revealed clustering of variants BA.2, JN.1, and BA.2.86, and distant separate cluster formed by XBB.1, thus supporting evolutionary divergence between JN.1 and XBB lineages [12]. In addition, lineages featuring the mutation F456L (including KP.2, KP.3, and XEC) [18], which has been shown to mediate substantial evasion of serum neutralization, were shown to comprise a cluster distinct from JN.1 (based on antigenic distance to D614G), regardless of the presence of other mutations [17]. In line with these data, neutralization assays performed on sera following immunization with monovalent XBB.1.5 vaccine show reduced neutralization titers against JN.1 compared with BA.2.86 [8], and significantly reduced neutralization of KP.2 compared with JN.1 [15], suggesting limited protection afforded by XBB.1.5 vaccines against antigenically distant sublineages [5].

The continual evolution of SARS-CoV-2 variants has resulted in a diverse landscape of JN.1-derived sublineages and recombinants [5], including the XEC strain, which is increasingly prevalent in Europe and North America [18]. Given the considerable antigenic divergence of the emerging lineages, individuals immunized only with the XBB.1.5 vaccine are likely to remain susceptible to infection caused by emerging SARS-CoV-2 variants, thus supporting the recommendation on updates to the antigenic composition of COVID-19 vaccines for the 2024-2025 season [5]. Numerous studies, both preclinical and clinical, have demonstrated the benefits of boosting immunization with variant-adapted mRNA-1273 vaccines in driving increased breadth of neutralization [41,42,44,45]. For example, boosting with the monovalent omicron-BA.1 matched mRNA-1273 vaccine (mRNA-1273.529) after a mRNA-1273 primary series in mice enhanced nAb responses against BA.1/BA.2 and, to a lesser degree, the historical SARS-CoV-2 D614G strain, and was associated with protection against BA.1 infection in the upper and lower respiratory tracts [44]. This has also been demonstrated in nonhuman primates, where boosting with either mRNA-1273 or mRNA-omicron vaccine following immunization with the primary series of mRNA-1273 expanded cross-reactive memory B cells and provided complete protection in the lungs after omicron challenge [45].

In conclusion, our results demonstrate the potency of the JN.1 and KP.2 variant vaccines in neutralizing the matched variants (JN.1, KP.2) and highlight the potential relevance of these vaccines in increasing neutralization of closely related subvariants such as KP.3, LA.2, and XEC. The lack of cross-neutralization between the XBB and JN.1 lineages highlights significant antigenic shifts that could re-emerge to drive immune escape, which, at present, is best mitigated by frequent boosting with variant-matched vaccines. The data presented here indicate that the currently licensed JN.1 and KP.2 vaccines are likely to continue to protect against the more recent emerging strains despite the steady antigenic drift from the vaccine strain. Continuous variant monitoring and testing will remain essential to identify the risk of immune escape and need for updated vaccines.

## Abbreviations

ELISA: enzyme-linked immunosorbent assay
GMT: geometric mean titer
JN.1: subvariant of omicron BA.2.86.1.1
KP.2: subvariant of omicron BA.2.86.1.1.11.1.2
LNP: lipid nanoparticle
NTD: N-terminal domain
RBD: receptor-binding domain
PBS: phosphate-buffer saline
PsVNA: pseudovirus neutralization assay
S-2P: spike protein with two proline substitutions within the heptad repeat 1 domain
VSV: vesicular stomatitis virus

## Acknowledgements

We gratefully acknowledge all data contributors, i.e., the Authors and their Originating laboratories responsible for obtaining the specimens, and their Submitting laboratories for generating the genetic sequence and metadata and sharing via the GISAID Initiative, on which some of this research is based. We thank Dr. Michael Whitt and Rita Kansal for support on pseudovirus production. Medical writing and editorial assistance were provided by Anja Varjačić, PhD, of MEDiSTRAVA in accordance with Good Publication Practice (GPP 2022) guidelines, funded by Moderna, Inc., and under the direction of the authors.

## Data Availability

All data supporting the findings of this study are available within this article and in the supplementary information.

## Ethics Approval

Animal experiments were carried out with the approval of the ModernaTX, Inc. Institutional Animal Care and Use Committee (IACUC Protocol 22-02-00) and in compliance with all applicable local and federal regulations, including the National Research Council’s Guide for the Care and Use of Laboratory Animals.

## Author Contributions

**Diana Wing Lee**: Data curation, Formal analysis, Investigation, Methodology, Visualization, Writing – original draft, Writing – review & editing. **Arshan Nasir**: Data curation, Formal analysis, Investigation, Methodology, Software, Visualization, Writing – review & editing. **Sayda Elbashir**: Conceptualization, Investigation, Resources, Supervision, Visualization, Writing – review & editing. **Hardik Jani**: Formal analysis, Investigation, Writing – review & editing. **Tessa Speidel**: Formal analysis, Writing – Review & editing. **Amy Gorrie**: Conceptualization, Formal analysis, Investigation, Writing – original draft, Writing – review & editing. **Daniela Montes Berrueta**: Formal analysis, Writing – review & editing. **Philippa Martin**: Formal analysis, Investigation, Writing – review & editing. **Swan Tan**: Data curation, Formal analysis, Writing – review & editing. **Jacob Hou**: Formal analysis, Visualization, Writing – original draft, Writing – review & editing. **Kath Hardcastle**: Conceptualization, Formal analysis, Investigation, Writing – original draft, Writing – review & editing. **Darin Edwards**: Conceptualization, Formal analysis, Investigation, Writing – original draft, Writing – review & editing. **Kai Wu**: Investigation, Methodology, Supervision, Writing – review & editing. **Andrea Carfi**: Conceptualization, Formal analysis, Writing – review & editing. **Yadunanda Budigi**: Conceptualization, Investigation, Methodology, Project administration, Supervision, Writing – original draft, Writing – review & editing.

