## Supplementary Materials for "mRNA-1273 vaccines adapted to JN.1 or KP.2 elicit cross-neutralizing responses against the JN.1 sublineages of SARS-CoV-2 in mice"

### Supplementary Methods

*Enzyme-linked immunosorbent assay (ELISA)*

Microtiter plates (96-well; Thermo Fisher Scientific) were coated with 100 µL of recombinant ancestral SARS-CoV-2 spike protein with 2 proline substitutions with the heptad repeat 1 domain (S-2P). After overnight incubation at 4°C, plates were blocked for 1.5 h at 4°C (SuperBlock, Thermo Fisher Scientific). Sera were serially diluted (5% goat serum in phosphate-buffered saline containing 0.1% Tween 20 [PBST]), added to plates, incubated for 2 hours at 37°C, and washed 3 times with PBST. Horseradish peroxidase-conjugated goat anti-mouse immunoglobulin G (IgG) (Southern Biotech) was diluted in 5% goat serum in PBST and added to the wells. Plates were incubated for 1 hour at 37°C, washed three times with PBST before the addition of 3,3',5,5'-tetramethylbenzidine (TMB) substrate (Thermo Fisher Scientific). Reactions were stopped by adding a TMB stop solution (Sera Care), and the optical density (OD) measurements were taken at 450 nm. Titers were determined using a 4-parameter logistic curve fit in Prism v.9 (GraphPad Software) and defined as the reciprocal dilution at approximately $OD_{450}$ = 1.0 (normalized to a mouse standard on each plate).

### Supplementary Figures

**Figure S1**. Binding antibody responses against prototype S-2P protein in BALB/c mice 14 days after primary series vaccination with mRNA-1273.167 (JN.1 variant vaccine; 1 µg) or mRNA-1273.712 (KP.2 variant vaccine; 1 µg) compared with mRNA-1273.815 (XBB.1.5 variant vaccine; 1 µg). **(A)** mRNA-1273.167; **(B)** mRNA-1273.712.

Data are shown as GMT ± 95% CI. GMT values are presented at the top of each figure, with red indicating the GMT for day 21 (3 weeks after dose 1) and blue indicating the GMT for Day 36 (2 weeks after dose 2). The dotted line indicates the LOD of 12.5 for the assay.

*Abbreviations:* CI, confidence interval; GMT, geometric mean titer; IgG, immunoglobulin G; LOD, limit of detection; PBS, phosphate-buffered saline; S-2P, spike protein with 2 proline substitutions within the heptad repeat 1 domain.

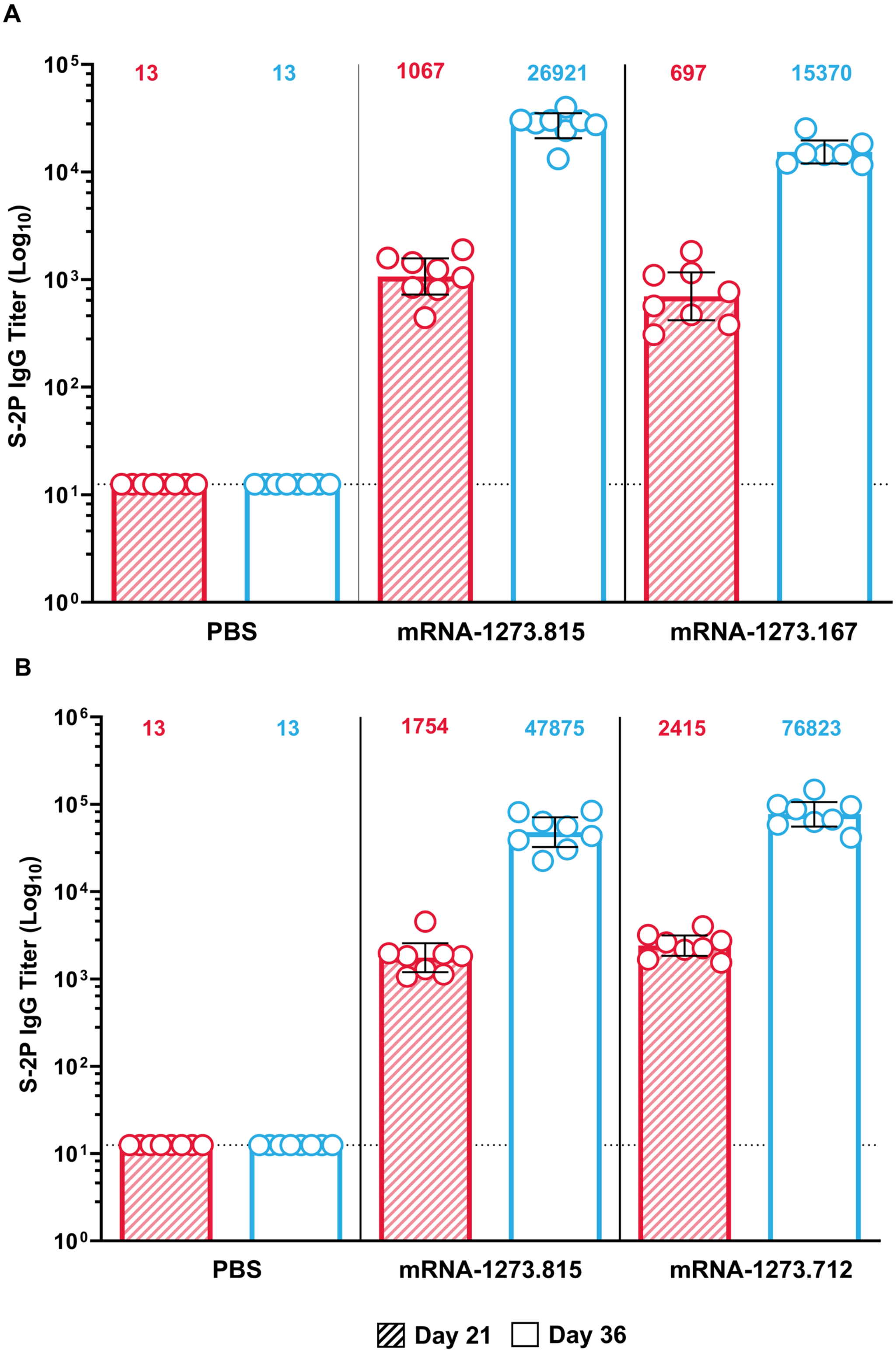
A
S-2P IgG Titer (Log10)
13
13
1067
26921
697
15370
PBS
mRNA-1273.815
mRNA-1273.167
B
S-2P IgG Titer (Log10)
13
13
1754
47875
2415
76823
PBS
mRNA-1273.815
mRNA-1273.712
Day 21
Day 36

**Figure S2.** Binding antibody responses against S-2P approximately 14 days following the administration of a single booster (third) dose in BALB/c mice. **(A)** mRNA-1273.167 (JN.1 variant vaccine; 1 μg), mRNA-1273.712 (KP.2 variant vaccine; 1 μg),[a] and mRNA-1273.815 (XBB.1.5 variant vaccine; 1 μg) administered ~19 days after immunization with 2-dose primary series of mRNA-1273 (ancestral strain). **(B)** mRNA-1273.167 and mRNA-1273.815 administered ~4 weeks after immunization with 2-dose primary series of mRNA-1273 (ancestral strain).

Data are shown as GMT ± 95% CI. GMT values are presented at the top of each figure, with red indicating pre-boost titers and blue indicating post-boost titers. The dotted line indicates the LOD of 12.5 for the assay.

[a]Serum samples were not available for ELISA testing from two mice that received KP.2 mRNA-1273 booster dose.

*Abbreviations:* CI, confidence interval; ELISA, enzyme-linked immunosorbent assay; GMT, geometric mean titer; IgG, immunoglobulin G; LOD, limit of detection; PBS, phosphate-buffered saline; S-2P, spike protein with 2 proline substitutions within the heptad repeat 1 domain.

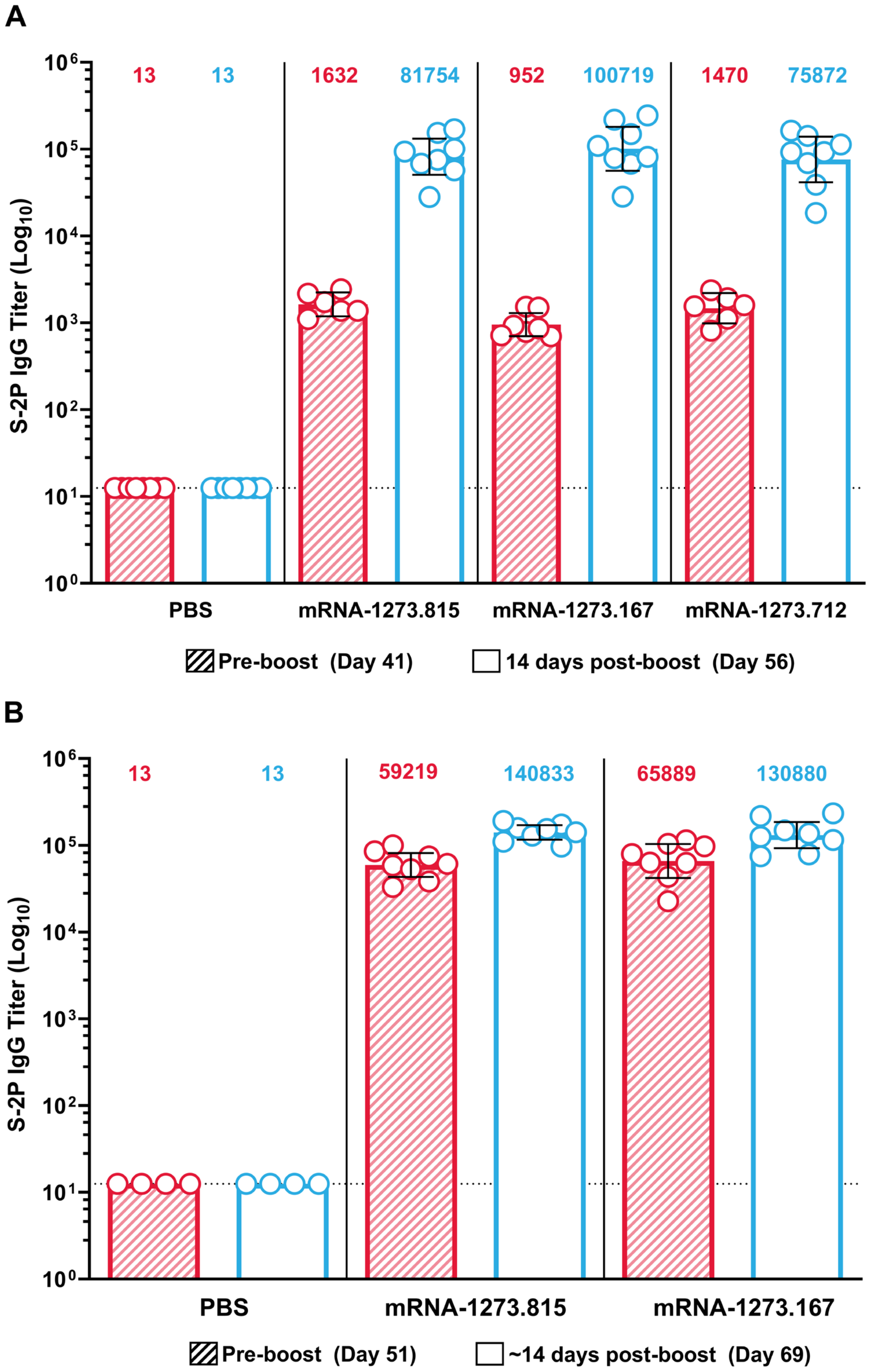
A
S-2P IgG Titer (Log10)
13
13
1632
81754
952
100719
1470
75872
PBS
mRNA-1273.815
mRNA-1273.167
mRNA-1273.712
Pre-boost (Day 41)
14 days post-boost (Day 56)
B
S-2P IgG Titer (Log10)
13
13
59219
140833
65889
130880
PBS
mRNA-1273.815
mRNA-1273.167
Pre-boost (Day 51)
~14 days post-boost (Day 69)
